# The future of artificial intelligence in digital pathology - results of a survey across stakeholder groups

**DOI:** 10.1101/2021.12.16.472990

**Authors:** Céline N. Heinz, Amelie Echle, Sebastian Foersch, Andrey Bychkov, Jakob Nikolas Kather

## Abstract

Artificial intelligence (AI) provides a powerful tool to extract information from digitized histopathology whole slide images. In the last five years, academic and commercial actors have developed new technical solutions for a diverse set of tasks, including tissue segmentation, cell detection, mutation prediction, prognostication and prediction of treatment response. In the light of limited overall resources, it is presently unclear for researchers, practitioners and policymakers which of these topics are stable enough for clinical use in the near future and which topics are still experimental, but worth investing time and effort into. To identify potentially promising applications of AI in pathology, we performed an anonymous online survey of 75 computational pathology domain experts from academia and industry. Participants enrolled in 2021 were queried about their subjective opinion on promising and appealing sub-fields of computational pathology with a focus on solid tumors. The results of this survey indicate that the prediction of treatment response directly from routine pathology slides is regarded as the most promising future application. This item was ranked highest in the overall analysis and in sub-groups by age and professional background. Furthermore, prediction of genetic alterations, gene expression and survival directly from routine pathology images scored consistently high across subgroups. Together, these data demonstrate a possible direction for the development of computational pathology systems in clinical, academic and industrial research in the near future.

## Introduction

Digital pathology is a “*dynamic, image-based environment that enables the acquisition, management and interpretation of pathology information generated from a digitized glass slide*” [1]. While today many pathology departments still operate in non-digital workflows, this is widely anticipated to change within the next five to ten years [2–5]. Digitalization of routine pathology workflows in itself can provide efficiency gains [6,7] while maintaining the quality of diagnosis [8]. However, the main benefit of digitizing pathology workflows is that it enables the application of new products, diagnostic systems and biomarkers, potentially expanding the information that can be extracted from tissue slides [9]. These new analytic approaches are referred to as “computational pathology” and include artificial intelligence (AI) approaches which are expected to add measurable value in the next decade [10]. Many of these AI methods are focused on cancer as a blueprint for the whole breadth of diagnostic pathology [5,11,12]. A recent systematic study categorized computational pathology in cancer applications in major groups (**Figure 1A**) [9]. One group contains basic applications aimed to automate tasks which are usually performed by pathologists, e.g. detection of tumor tissue [13], subtyping of tumors [14] and automating grading [15,16]. Another major group is that of advanced applications, i.e. extraction of information from slides beyond current clinical routine. In particular, these applications include three categories. First, the prediction of genetic and molecular alterations directly from digitized hematoxylin and eosin (H&E)-stained slides, such as mutations [17–19], gene expression [20], microsatellite instability [21,22] and epigenetic features; second, prediction of survival directly from H&E images [23,24]; and lastly the direct end-to-end prediction of response to a particular treatment in cancer, including immunotherapy [25] and targeted therapy [26,27]. In addition to these “basic” and “advanced” approaches, another broad category of computational pathology focuses on “enabling tasks”, such as quality control [28], detection of cells or tissue structures in images (for the purpose of subsequent quantification [29,30]) or quantification of immunohistochemical (IHC) stains [31].

**Figure 1:**
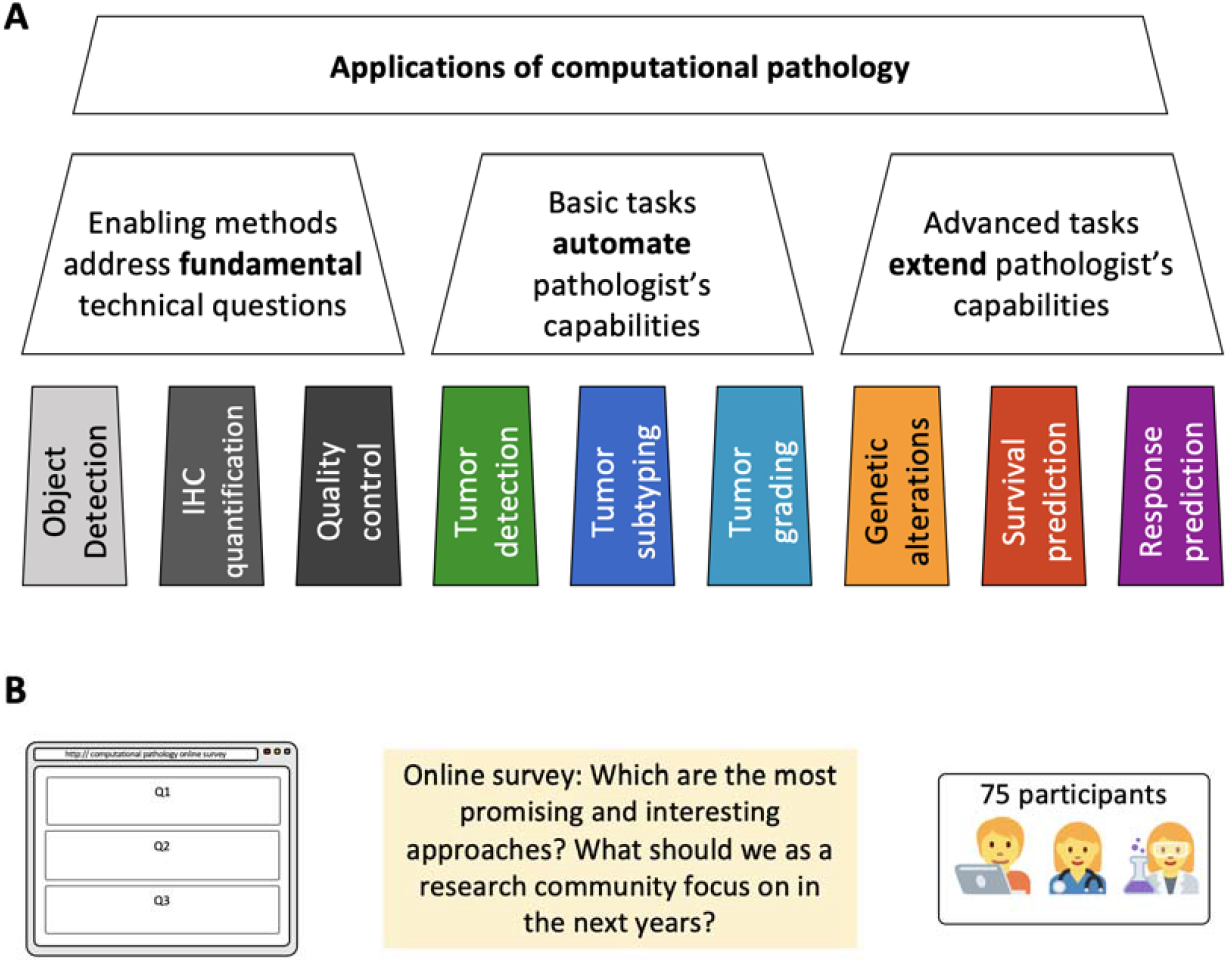
Applications of computational pathology and setup of this survey. (**A**) Based on Echle et al.[9], applications of computational pathology can be categorized as “basic” or “advanced”. In this study, enabling technologies were introduced as an additional category. (**B**) We performed an online survey among 75 participants who are diverse stakeholders in the field. Participants were asked to report their subjective experience or opinion.

However, it is currently unclear which of these approaches are most likely to bring tangible benefits in the near future. Therefore, we designed and performed an online survey among 75 stakeholders from different professional areas dealing with computational pathology. We focused on applications of AI in the analysis of images derived from solid tumors. We aimed to systematically investigate which computational pathology applications are considered as being most “promising”, “interesting” and “worth to focus” on. Here, we present, quantify and interpret the results of this systematic survey.

## Methods

### Ethics statement

We performed an anonymous online survey among professionals working in the field of computational pathology. All participants voluntarily participated in this study and no benefit or disadvantage was associated with participation or non-participation in this survey. The identity of the participants is unknown to the study authors. No patient data were used for this study. The protocol of this study is in accordance with the Declaration of Helsinki.

### Survey design

The survey was split in two parts: First, participants reported their demographic and professional background in six questions (**Table 1, “First, some questions on your background”**). Levels of experience were subdivided into three larger categories, covering aspects of previous involvement in AI-based image analysis projects, hands-on experience in computer programming and participants’ own training of an artificial neural network. Second, for each application category, a numerical rating scale was used to indicate the priority participants assigned to a given category (**Table 1, “Which applications of AI in digital pathology do you personally find promising or interesting? - AI for…”**). Categories were taken from Echle et al. [9], including “basic” and “advanced” tasks. In addition, we added two categories (object detection and IHC quantification) to compare the end-to-end histopathology approach to conventional digital pathology approaches. Lastly, to give participants the opportunity to make additional comments, the survey included two optional free-text questions (**Table 1,** “**Additional free text fields”**). All response options were either categorical, a numerical rating scale or free text responses (**Table 1**).

**Table 1:** Questionnaire description and reference.

| Item # | Survey Question | Response options | Explanation / Reference |
| --- | --- | --- | --- |
|  | **"First, some questions on your background"** |  |  |
| 1 | How did you come across this survey? | Categorical, select one of the following:<br>1. An online course ("SFP AI in pathology course", "EMPAIA Academy", ...)<br>2. A talk at a scientific conference ("DGP conference", "ECP", ...)<br>3. Social media (Twitter, LinkedIn, Facebook)<br>4. Personal connections<br>5. Other | Basic information about the professional and biographic background |
| 2 | You were born between... | Categorical, select one of the following:<br>1. 1959 or earlier<br>2. 1960 - 1969<br>3. 1970 - 1979<br>4. 1980 - 1989<br>5. 1990 - 1999<br>6. 2000 or later |  |
| 3 | What best describes your background? | Categorical, select one of the following:<br>1. Pathology<br>2. Oncology / Clinical<br>3. Medical / Translational research<br>4. Technical (Engineering, Computer Science) research<br>5. Industry<br>6. Other |  |
| 4 | Have you ever been involved in AI based image analysis projects? | Categorical, select one of the following:<br>1. Yes<br>2. No<br>3. Not sure | Information about the previous experience of participants |
| 5 | Do you have hands-on experience in computer programming? | Categorical, select one of the following:<br>1. Yes<br>2. No<br>3. Not sure |  |
| 6 | Have you ever trained an artificial neural network? | Categorical, select one of the following:<br>1. Yes<br>2. No<br>3. Not sure |  |
|  | “Which applications of AI in digital pathology do you personally find promising or interesting? - AI for...” |  |  |
| 7 | detection of specific cell types in H&E images (e.g. lymphocytes) | Rating scale from 1 (not interesting) to 10 (very interesting) | New category “Object detection” |
| 8 | detection of large structures in H&E images (e.g. nerves, blood vessels, lymph follicles, ...) |  |  |
| 9 | classifying tissue types in H&E images (e.g. epithelium, stroma, muscle, fat ...) |  | Echle [9]: “basic / tumor detection” |
| 10 | quantification of immunostains in histology images (e.g. Her2, Ki67, PDL1, ...) |  | New category “immunostains” |
| 11 | prediction of genetic mutations from H&E images of tumors (e.g. BRAF mutation, EGFR mutation ...) |  | Echle [9]: “advanced / prediction of genetic alterations” |
| 12 | prediction of gene expression from H&E images of tumors |  |  |
| 13 | prediction of epigenetic features from H&E images of tumors (e.g. methylation, ...) |  |  |
| 14 | automatic tumor grading in H&E images (e.g. Prostate cancer Gleason grading, or grading of differentiation in colorectal cancer) |  | Echle [9]: “basic / grading” |
| 15 | subtyping tumor types based on morphology (e.g. adenocarcinoma vs. squamous cell carcinoma in lung cancer) |  | Echle [9]: “basic / subtyping” |
| 16 | prediction of survival directly from H&E images of solid tumors |  | Echle [9]: “advanced / survival prediction” |
| 17 | prediction of tissue and image quality directly from H&E images |  | New category “quality control” |
| 18 | prediction of treatment response directly from H&E images of solid tumors |  | Echle [9]: “advanced / response prediction” |
|  | Additional free text fields |  |  |
| 19 | Which other applications of AI in computational pathology would you find promising or interesting? | Free text, optional | receive additional ideas which the pre-specified categories cannot capture |
| 20 | In your opinion, how can the use of AI in medical research be further improved? | Free text, optional |  |

### Implementation and distribution of the survey

The survey was performed using Google Forms (Google Inc., Mountain View, CA, https://www.google.com/forms/about/). The survey was active from 3rd of June 2021 until 20th of July 2021 and was closed when 75 responses were recorded (pre-defined stopping criterion). The survey was advertised during online events (“Artificial Intelligence in Pathology Course” sponsored by the French Society for Pathology in June 2021 and lectures at scientific symposia in June 2021). In addition, the survey was promoted on social networks Twitter and LinkedIn via professional accounts of the authors.

### Data analysis and data availability

For analysis of the data, responses on a numerical rating scale were primarily analyzed by median and interquartile range (IQR). To visualize differences between groups in an exploratory analysis, the mean and standard deviation (SD) was used. To test whether responses in a given category were significantly higher than in another category, Kruskal-Wallis Test was used. No correction for multiple testing was applied and only p-values below 0.01 were considered significant. Correlation between categories was assessed via Spearman’s correlation analysis. Statistical analyses were carried out using Microsoft Excel and Python/SciPy. Analyses were carried out for all participants (full cohort) and pre-defined subgroups (older vs. younger participants and medical vs. non-medical background). Younger participants (n = 54) were defined as those born between 1980 and 1999 while older (n = 21) participants were those born between 1960 and 1979. None of the 75 total respondents (full cohort) were born 1959 or earlier, nor 2000 or later. Based on the professional background, a “medical” subgroup (n = 36) covered the proposed disciplines “Pathology”, “Oncology/Clinical” and “Medical/Translational research”, while the “nonmedical” subgroup (n = 39) included all other participants, i.e. those from fields of “Technical (Engineering, Computer Science) research”, “Industry” and “Other”. Anonymized raw data are available in **Suppl. Table 1**.

## Results

### Basic demographic and professional background of participants

In June and July 2021, we performed an anonymous online survey among self-identified members from the field of computational pathology (**Figure 1B, Suppl. Table 2**). 56 (75%) participants were born before and 19 (25%) participants were born after 1990 (**Figure 2A**, **Suppl. Table 3**). 36 (48%) participants had a medical background while 39 (52%) participants had a technical background (**Figure 2B**). 43 (57%) participants accessed the survey through social media (Twitter, LinkedIn, Facebook), where the survey was shared. 14 (19%) participants linked to personal connections, 13 (17%) individuals came across the survey form by participating in an online course, 4 (5%) through a talk at a scientific conference and 1 (1%) participant through another source (**Figure 2C**). Regarding previous experience with AI in computational pathology, 49 (65%) participants had hands-on-experience in computer programming (**Figure 2D**), 42 (56%) participants reported having trained a neural network before (**Figure 2E**) and 58 (77%) participants reported having been involved in computer-based image analysis projects (**Figure 2F**).

**Figure 2:**
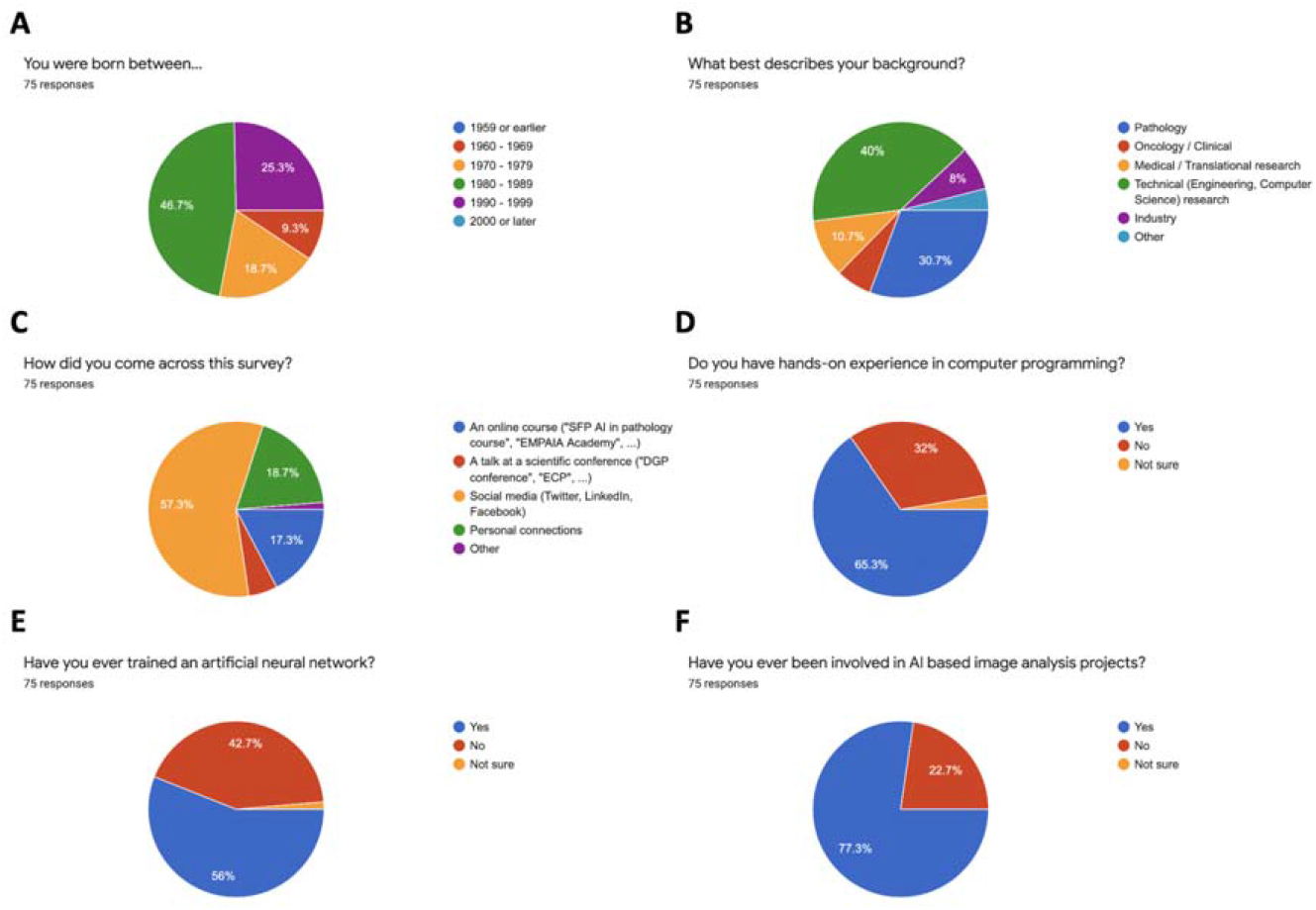
Basic demographics and previous experience of survey participants. (**A**) Self-reported age among participants, (**B**) Professional background of participants. (**C**) Distribution of the survey (**D**) Computer programming experience among participants, (**E**) Hands-on machine learning experience among participants, (**F**) project experience of participants.

### Which applications are most promising overall?

Participants were asked to rank different applications of AI in digital pathology on a numerical rating scale from 1 to 10, where 1 represents the least important and 10 the most interesting and/or promising application in their subjective opinion. According to all participants, the most promising application for AI in digital pathology was predicting treatment response directly from H&E images of solid tumors, reaching an overall arithmetic mean of 9.09 (median of 10, **Figure 3A**). In a pairwise comparison of categories, prediction of treatment response was scored significantly (p < 0.01) higher than all items in “enabling” or “basic” categories (**Suppl. Figure 1A**). Also, predicting genetic mutations as well as gene expression directly from H&E image were considered second and third most interesting with an almost equal score of mean 8.64 (median of 10) and mean 8.63 (median of 10), respectively (**Figure 3A**). Taken together, these data show that “advanced” tasks according to Echle et al. [9] (**Figure 1A**) reached the highest scores among the participants. In contrast, “enabling” technologies scored lower: With an arithmetic mean of 6.83 (median of 8), AI for detection of large structures in H&E images was considered to be the overall least promising application of AI in digital pathology and the scores were significantly (p<0.01) lower than all scores of items in the “advanced” category (**Figure 3**). In addition, we analyzed the correlation between responses, i.e. which categories were usually rated in a similar way by participants. This analysis showed that individual items within the categories “enabling technologies”, “automation” and “advanced” (**Figure 1A**) generally clustered together, i.e. were often rated in a similar way by the participants (**Suppl. Figure 1B**).

**Figure 3:**
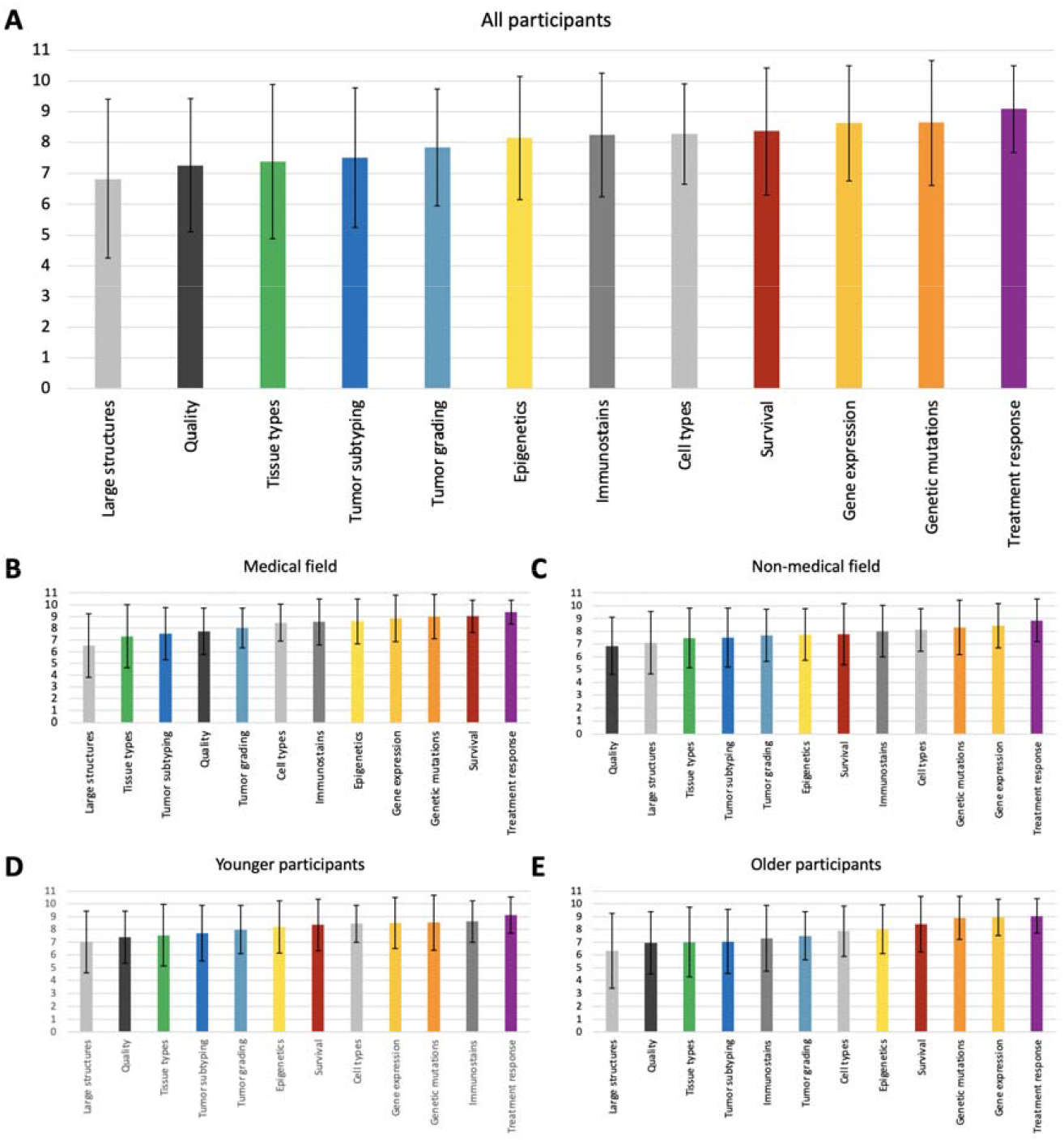
Ranking of responses in all participants and subgroups. (**A**) Arithmetic mean and standard deviation for all participants. (**B**) For participants with a medical professional background. (**C**) For participants with a non-medical professional background. (**D**) Younger participants born between 1980 and 1999. (**E**) Older participants born between 1960 and 1979.

### Subgroup analysis by professional and demographic background

Next, we assessed the influence of professional background and age on participants’ responses. Among participants with a professional background in a medical field as well as from a non-medical field (**Figure 2B**), as well as for younger and older participants (**Suppl. Table 3**), prediction of treatment response was consistently ranked highest. Detection of large structures in histopathology images was consistently ranked lowest or second-to-lowest. In all four subgroups, prediction of survival was among the top 50% ranked targets and prediction of molecular features from H&E images was also consistently ranked high. Interestingly, prediction of survival was ranked higher by participants with a medical (**Figure 3B**) than with a non-medical (**Figure 3C, Suppl. Table 4**) background. Another interesting finding related to the quantification of immunostains which was ranked second-highest by the younger subgroup (**Figure 3D, Suppl. Table 5**). Apart from these observations, no striking deviation from the overall ranking results (**Figure 3A**) was made for any of the subgroups.

### Analysis of free text responses

In addition to the quantitative analyses on numerical rating scales, participants were asked for free-text suggestions on application of AI in computational pathology. A total of 32 answers were given to the first open question. Most participants (n = 8) suggested to use AI for time consuming and laborious routine tasks in the pathological field (e.g. virtual staining and automated measurement) to integrate AI based procedures into clinical practice and improve the efficiency of pathologists and medical researchers in their daily workflow (**Figure 4A, Suppl. Table 6**). Another common response (n = 6) was to use AI for quality control purposes, especially quality of the tissue, quality of staining and quality of scans. Furthermore, respondents showed their interest in using AI for detection of subtle morphological features directly from digital histological images such as a more detailed cell and tissue characterization (e.g. in shape and structure) (n = 3) or for detection of bacteria (e.g. acid-fast bacteria (AFB) or H. pylori) (n = 2).

**Figure 4:**
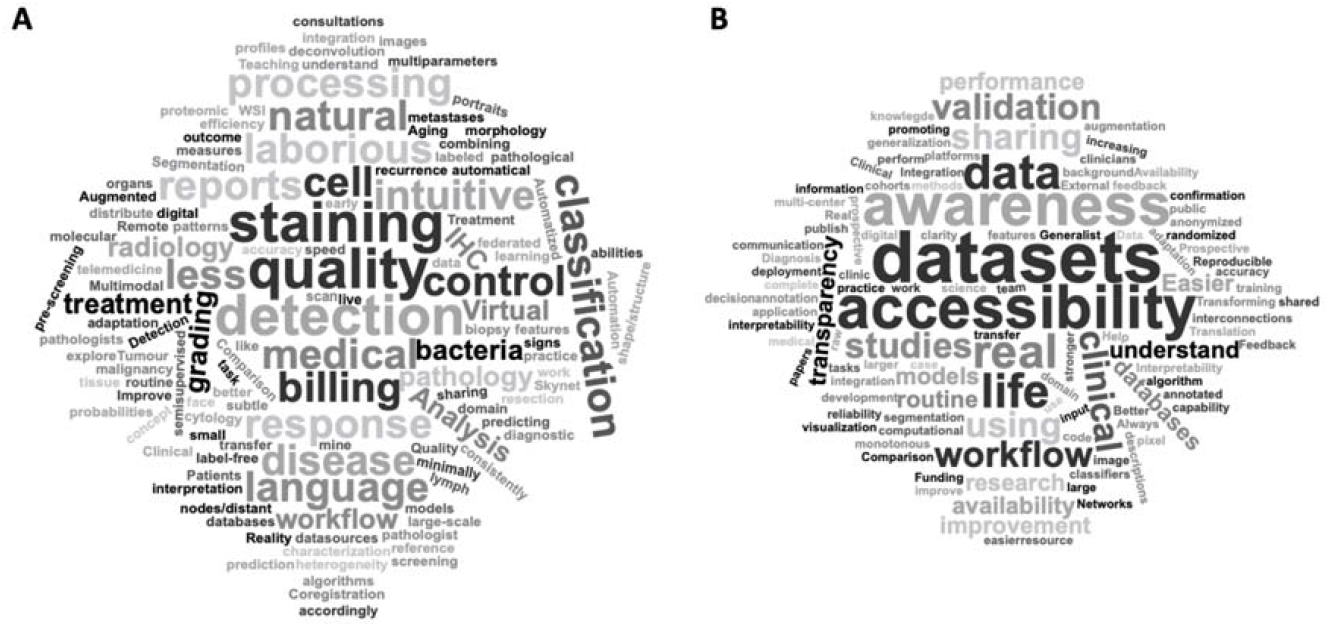
Qualitative analysis of responses to free text questions. **(A)** ‘Which other applications of AI in computational pathology would you find promising or interesting?’, **(B)** ‘In your opinion, how can the use of AI in medical research be further improved?’

In a second free-text question, participants were asked for suggestions how AI could improve medical research in general (**Figure 4B, Suppl. Table 7**). A total of 36 answers were given to this question. The most common (n = 13) suggestion was to reinforce integration of AI into clinical real-life settings to improve routine medical and pathological tasks in a durable way. Improving availability of large data sets, clinical cohorts and established AI codes as well as making the latter publicly accessible came as a close second (n = 11). The importance of comparing AI mined results with the real existing clinical data (e.g. by augmenting feedback from pathologists or clinicians) and thereby working towards increasing reproducibility of obtained results was another common suggestion (n = 9). Six respondents suggested interpretability of AI models for medical professionals as an important area, arguing that this could create stronger interconnections between technical and non-technical fields in order to create transparency for AI-based medical image analysis. Equally represented was the implementation of prospective and multi-center studies in the context of AI-based image analysis in medical research fields to start integrating AI workflows into clinical practice. Lastly, four participants explicitly wished for more public awareness regarding the overall existence of AI in a medical research context.

## Discussion

### Key points and comparison to previous work

We report the results of an online survey among various stakeholders in computational pathology which aimed to identify promising applications of AI in computational pathology. Our quantitative and qualitative analysis aimed to identify research areas of interest which could help stakeholders to sensibly allocate resources in academic and industry settings. Based on previous work [9], we divided the fields of application into enabling technologies, automation of pathology workflows and advanced tasks (**Figure 1**). We found that advanced applications of AI were consistently rated as most interesting across all subgroups of participants (**Figure 3**). Responses in free-text questions pointed out the need of using AI to automate time-consuming diagnostic tasks and real-world validation of pathology AI systems. This is in line with recent initiatives to increase the level of evidence of computational pathology systems [32]. Another prominent point was improvement of data availability and data sharing which is in line with recent approaches to standardize reporting of dataset-related features in AI biomarker studies [33]. Compared to previous surveys about AI in digital pathology [34–36], our study did not focus on implementation and training, but rather on the development of new AI approaches for digital pathology.

### Limitations

A limitation of our method is a possible non-response bias. Participants in the survey were recruited by disseminating invitations at professional conferences and on social networks targeting a particular audience. The decision of individuals in the target population to participate in the survey could bias the results [37,38]. Pathology experts not familiar with or even critical to digital pathology and AI applications might have a different opinion on what the research should focus on in this regard. Another limitation is that the survey was restricted to English-speaking participants, although no additional data was collected to investigate the geographical distribution of participants. Finally, this survey provides just a snapshot in time and topics of subjective relevance could change over time. In this sense, our survey may serve as a starting point for the follow-up studies over a certain time period.

### Outlook

As of 2021, academic prototypes of artificial intelligence (AI) systems are available in many domains of computational pathology [9]. Implementation is mostly limited by organizational problems, such as running validation studies, digitization of diagnostic workflows or collection of samples to train AI systems [39–41]. In this light, we anticipate that in the next ten years, a focus for researchers in academia and industry will be on the development of new computational pathology solutions. Such solutions could run within an existing digital pathology environment, but it is currently unclear which applications would provide the most benefit to stakeholders, including patients. This survey focused on the perception of promising applications of AI in digital pathology. The results of our survey indicate that advanced tasks, i.e. predicting features from pathology images which extend pathologists’ capabilities, is consistently judged as the most relevant application. In addition, our study demonstrates that a survey can yield quantifiable insight about future directions of a translational research field, which merits further validation in large-scale surveys.

## Supplementary Data

**Suppl. Table 1:** All raw data used in this analysis. [external file]

**Suppl. Table 2:** Median (med) and interquartile range (IQR) for all subgroups for all questionnaire items. Questionnaire items were judged on a scale from 1 to 10, 10 indicating highest agreement / importance.

| Item # | Questionnaire Item | Overall N=75 | Young N=54 | Older N=21 | Medical N=23 | Other N=52 |
| --- | --- | --- | --- | --- | --- | --- |
| 7 | detection of specific cell types in H&E images (e.g. lymphocytes) | med=8<br>IQR=2,5 | med=8.5<br>IQR=2 | med=8<br>IQR=3 | med=9<br>IQR=2 | med=8<br>IQR=3 |
| 8 | detection of large structures in H&E images (e.g. nerves, blood vessels, lymph follicles, ...) | med=8<br>IQR=3.5 | med=8<br>IQR=2.75 | med=7<br>IQR=5 | med=7<br>IQR=3 | med=8<br>IQR=3 |
| 9 | classifying tissue types in H&E images (e.g. epithelium, stroma, muscle, fat ...) | med=8<br>IQR=4 | med=8<br>IQR=4 | med=8<br>IQR=4 | med=8<br>IQR=4.25 | med=8<br>IQR=3.50 |
| 10 | quantification of immunostains in histology images (e.g. Her2, Ki67, PDL1, ...) | med=9<br>IQR=3 | med=9<br>IQR=2 | med=8<br>IQR=5 | med=9.5<br>IQR=2 | med=8<br>IQR=3 |
| 11 | prediction of genetic mutations from H&E images of tumors (e.g. BRAF mutation, EGFR mutation ...) | med=10<br>IQR=2 | med=10<br>IQR=2 | med=10<br>IQR=2 | med=10<br>IQR=1.25 | med=9<br>IQR=2.50 |
| 12 | prediction of gene expression from H&E images of tumors | med=10<br>IQR=2 | med=9,5<br>IQR=2,75 | med=10<br>IQR=2 | med=10<br>IQR=2 | med=9<br>IQR=3 |
| 13 | prediction of epigenetic features from H&E images of tumors (e.g. methylation, ...) | med=8<br>IQR=3 | med=8<br>IQR=3 | med=8<br>IQR=3 | med=9<br>IQR=2 | med=8<br>IQR=4 |
| 14 | automatic tumor grading in H&E images (e.g. Prostate cancer Gleason grading, or grading of differentiation in colorectal cancer) | med=8<br>IQR=3,5 | med=8<br>IQR=3 | med=8<br>IQR=3 | med=8<br>IQR=4 | med=8<br>IQR=2 |
| 15 | subtyping tumor types based on morphology (e.g. adenocarcinoma vs. squamous cell carcinoma in lung cancer) | med=8<br>IQR=3 | med=8<br>IQR=2.75 | med=7<br>IQR=3 | med=8<br>IQR=3.25 | med=8<br>IQR=3 |
| 16 | prediction of survival directly from H&E images of solid tumors | med=9<br>IQR=3 | med=9<br>IQR=2.75 | med=9<br>IQR=3 | med=10<br>IQR=2 | med=8<br>IQR=3.50 |
| 17 | prediction of tissue and image quality directly from H&E images | med=8<br>IQR=3 | med=8<br>IQR=3 | med=8<br>IQR=3 | med=8<br>IQR=4 | med=7<br>IQR=2 |
| 18 | prediction of treatment response directly from H&E images of solid tumors | med=10<br>IQR=2 | med=10<br>IQR=2 | med=10<br>IQR=2 | med=10<br>IQR=1.25 | med=9<br>IQR=2 |

**Suppl. Table 3:** Precise representation of the individual age subgroups defined by a ten-year range of birth years respectively. Birth years are listed in ascending order.

| Year of birth | Overall percentage | Number of participants |
| --- | --- | --- |
| 1960 - 1969 | 9.30% | N=7 |
| 1970 - 1979 | 18.70% | N=14 |
| 1980 - 1989 | 46.70% | N=35 |
| 1990 - 1999 | 25.30% | N=19 |

**Suppl. Table 4:** Detailed overview of the professional background subgroup analysis including the two secondary established subgroups “medical” and “non-medical” and showing arithmetic mean, median and interquartile range (IQR) for the top three ranked applications respectively.

| Field | Most relevant applications |
| --- | --- |
| Medical | 1. AI for prediction of treatment response<br>$\bar{x} = 9.36/10$; median: 10; IQR: 1.25<br>2. AI for prediction of survival<br>$\bar{x} = 9.03/10$; median: 10; IQR: 2<br>3. AI for prediction of genetic mutations<br>$\bar{x} = 9.00/10$; median: 10; IQR: 1.25 |
| Non-medical | 1. AI for prediction of treatment response<br>$\bar{x} = 8.85/10$; median: 9; IQR: 2<br>2. AI for prediction of gene expression<br>$\bar{x} = 8.44/10$; median: 9; IQR: 3<br>3. AI for prediction of genetic mutations<br>$\bar{x} = 8.31/10$; median: 9; IQR: 2.50 |

**Suppl. Table 5:**
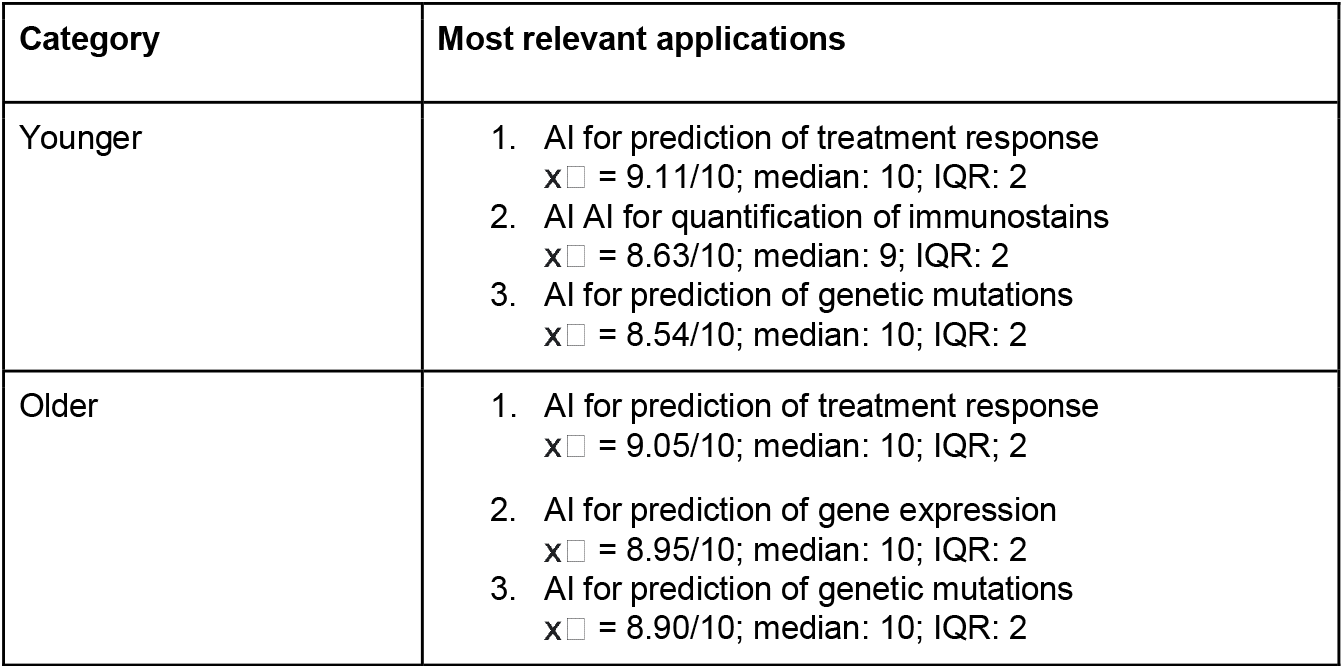
Detailed overview of age subgroup analysis including the two secondary established age subgroups “younger” and “older” and showing arithmetic mean, median and interquartile range (IQR) for the top three ranked applications respectively.

**Suppl. Table 6:** Overview of respondents’ most frequently given answers for open question number 1 (‘Which other applications of AI in computational pathology would you find promising or interesting?’). Different answer categories were established during free text question analysis and are listed in decreasing order.

| Answer category | Number of repetitions | Total answers |
| --- | --- | --- |
| Using AI for automation of time consuming routine tasks (e.g. virtual staining) | 9 | 32 |
| Using AI for quality control of tissue and scans | 6 | 32 |
| Using AI for detection of subtle features from H&E images (e.g. more detailed cell characterization) | 4 | 32 |
| Using AI for prediction of treatment response directly from digitized slides | 3 | 32 |
| Using AI for detection of bacteria | 2 | 32 |
| Using AI for teaching | 1 | 32 |

**Suppl. Table 7:**
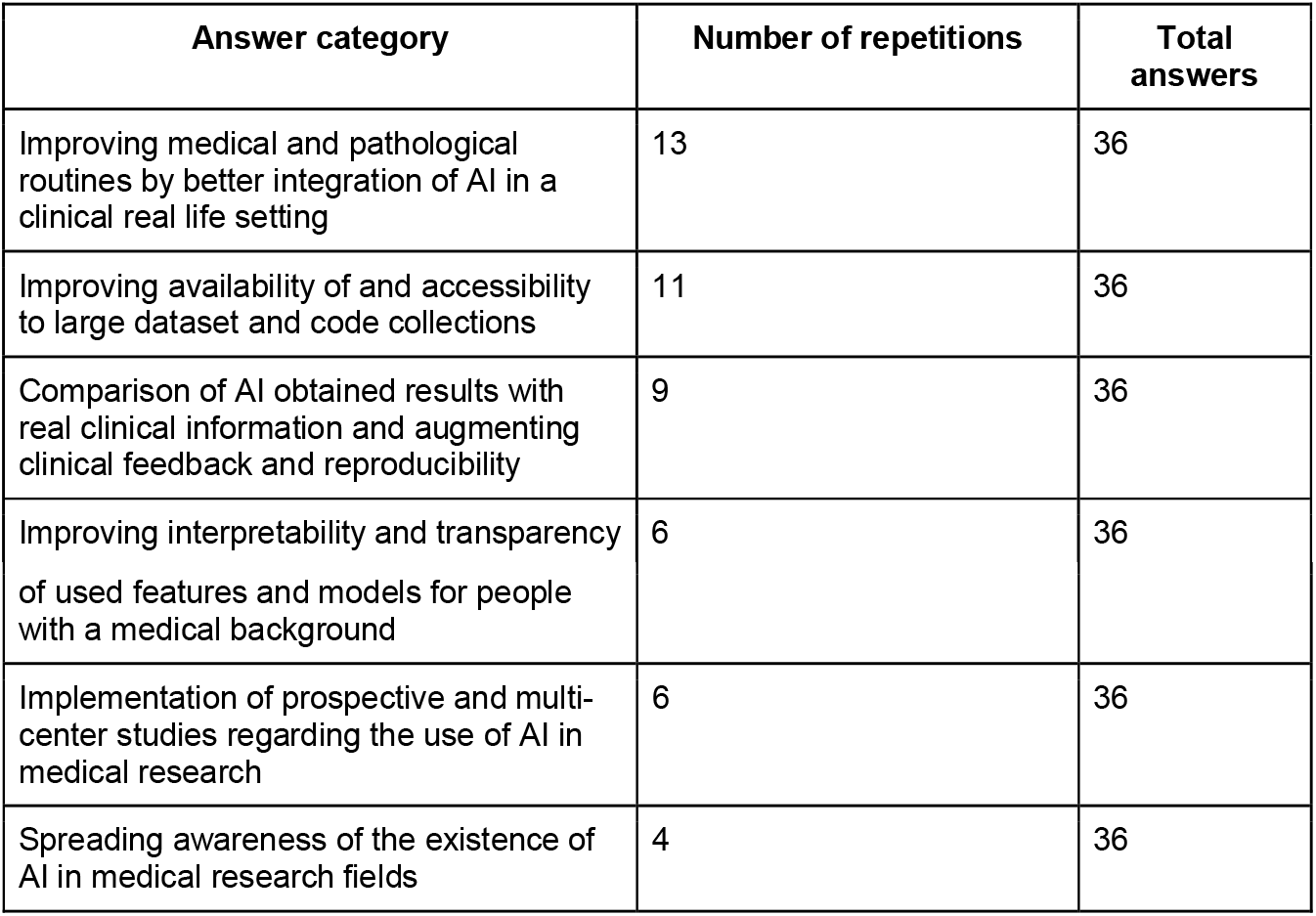
Overview of respondents’ most frequently given answers for open question number 2 (‘In your opinion, how can the use of AI in medical research be further improved?’). Different answer categories were established during free text question analysis and are listed in decreasing order.

**Suppl. Figure 1:**
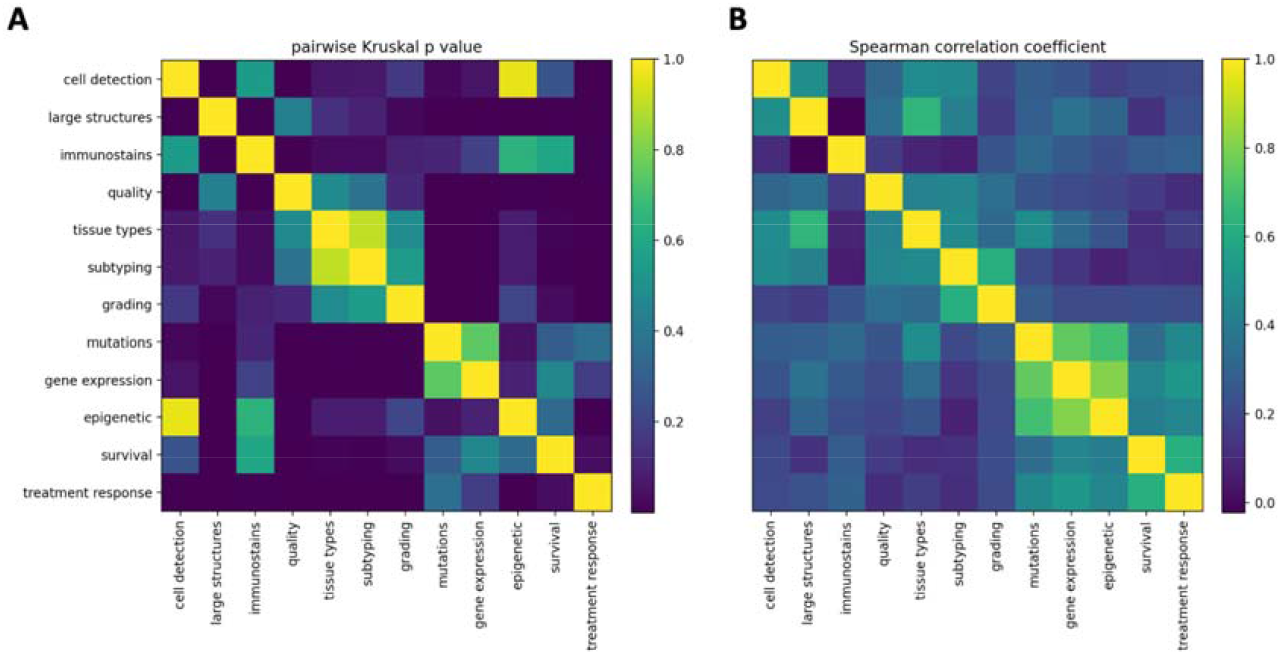
(A) Statistical significance of pairwise differences between scores in categories (pairwise Kruskal p value), (B) Correlation between scores in categories (Spearman correlation coefficient) in N=75 participants.

## Author contributions

CNH, AE and JNK designed the survey. CNH collected the data and performed the analyses. CNH wrote the manuscript and all authors contributed to editing the final version. All authors contributed to the interpretation of results and collectively made the decision to submit for publication.

## References

1. About Digital Pathology. [cited 16 Dec 2021]. Available: https://digitalpathologyassociation.org/about-digital-pathology

2. Wang S, Yang DM, Rong R, Zhan X, Xiao G. Pathology Image Analysis Using Segmentation Deep Learning Algorithms. Am J Pathol. 2019;189: 1686–1698.

3. Barsoum I, Tawedrous E, Faragalla H, Yousef GM. Histo-genomics: digital pathology at the forefront of precision medicine. Diagnosis (Berl). 2019;6: 203–212.

4. Jiang F, Jiang Y, Zhi H, Dong Y, Li H, Ma S, et al. Artificial intelligence in healthcare: past, present and future. Stroke Vasc Neurol. 2017;2: 230–243.

5. Sultan AS, Elgharib MA, Tavares T, Jessri M, Basile JR. The use of artificial intelligence, machine learning and deep learning in oncologic histopathology. J Oral Pathol Med. 2020;49: 849–856.

6. Hanna MG, Reuter VE, Hameed MR, Tan LK, Chiang S, Sigel C, et al. Whole slide imaging equivalency and efficiency study: experience at a large academic center. Mod Pathol. 2019;32: 916–928.

7. Vodovnik A. Diagnostic time in digital pathology: A comparative study on 400 cases. J Pathol Inform. 2016;7: 4.

8. Pantanowitz L, Sinard JH, Henricks WH, Fatheree LA, Carter AB, Contis L, et al. Validating whole slide imaging for diagnostic purposes in pathology: guideline from the College of American Pathologists Pathology and Laboratory Quality Center. Arch Pathol Lab Med. 2013;137: 1710–1722.

9. Echle A, Rindtorff NT, Brinker TJ, Luedde T, Pearson AT, Kather JN. Deep learning in cancer pathology: a new generation of clinical biomarkers. British Journal of Cancer. 2020. doi:10.1038/s41416-020-01122-x

10. Kather JN, Calderaro J. Development of AI-based pathology biomarkers in gastrointestinal and liver cancer. Nat Rev Gastroenterol Hepatol. 2020;17: 591–592.

11. Noorbakhsh J, Farahmand S, Foroughi Pour A, Namburi S, Caruana D, Rimm D, et al. Deep learning-based cross-classifications reveal conserved spatial behaviors within tumor histological images. Nat Commun. 2020;11: 6367.

12. Litjens G, Kooi T, Bejnordi BE, Setio AAA, Ciompi F, Ghafoorian M, et al. A survey on deep learning in medical image analysis. Med Image Anal. 2017;42: 60–88.

13. Campanella G, Hanna MG, Geneslaw L, Miraflor A, Werneck Krauss Silva V, Busam KJ, et al. Clinical-grade computational pathology using weakly supervised deep learning on whole slide images. Nat Med. 2019;25: 1301–1309.

14. Lu MY, Williamson DFK, Chen TY, Chen RJ, Barbieri M, Mahmood F. Data-efficient and weakly supervised computational pathology on whole-slide images. Nature Biomedical Engineering. 2021; 1–16.

15. Ström P, Kartasalo K, Olsson H, Solorzano L, Delahunt B, Berney DM, et al. Artificial intelligence for diagnosis and grading of prostate cancer in biopsies: a populationbased, diagnostic study. Lancet Oncol. 2020;21: 222–232.

16. Bulten W, Pinckaers H, van Boven H, Vink R, de Bel T, van Ginneken B, et al. Automated deep-learning system for Gleason grading of prostate cancer using biopsies: a diagnostic study. Lancet Oncol. 2020;21: 233–241.

17. Kather JN, Heij LR, Grabsch HI, Loeffler C, Echle A, Muti HS, et al. Pan-cancer imagebased detection of clinically actionable genetic alterations. Nature Cancer. 2020;1: 789–799.

18. Fu Y, Jung AW, Torne RV, Gonzalez S, Vöhringer H, Shmatko A, et al. Pan-cancer computational histopathology reveals mutations, tumor composition and prognosis. Nature Cancer. 2020; 1–11.

19. Loeffler CML, Ortiz Bruechle N, Jung M, Seillier L, Rose M, Laleh NG, et al. Artificial Intelligence–based Detection of FGFR3 Mutational Status Directly from Routine Histology in Bladder Cancer: A Possible Preselection for Molecular Testing? European Urology Focus. 2021. doi:10.1016/j.euf.2021.04.007

20. Schmauch B, Romagnoni A, Pronier E, Saillard C, Maillé P, Calderaro J, et al. A deep learning model to predict RNA-Seq expression of tumours from whole slide images. Nature Communications. 2020. doi:10.1038/s41467-020-17678-4

21. Muti HS, Heij LR, Keller G, Kohlruss M, Langer R, Dislich B, et al. Development and validation of deep learning classifiers to detect Epstein-Barr virus and microsatellite instability status in gastric cancer: a retrospective multicentre cohort study. The Lancet Digital Health. 2021;0. doi:10.1016/S2589-7500(21)00133-3

22. Echle A, Grabsch HI, Quirke P, van den Brandt PA, West NP, Hutchins GGA, et al. Clinical-Grade Detection of Microsatellite Instability in Colorectal Tumors by Deep Learning. Gastroenterology. 2020;159: 1406–1416.e11.

23. Skrede O-J, De Raedt S, Kleppe A, Hveem TS, Liestøl K, Maddison J, et al. Deep learning for prediction of colorectal cancer outcome: a discovery and validation study. Lancet. 2020;395: 350–360.

24. Kather JN, Krisam J, Charoentong P, Luedde T, Herpel E, Weis C-A, et al. Predicting survival from colorectal cancer histology slides using deep learning: A retrospective multicenter study. PLoS Med. 2019;16: e1002730.

25. Johannet P, Coudray N, Donnelly DM, Jour G, Illa-Bochaca I, Xia Y, et al. Using Machine Learning Algorithms to Predict Immunotherapy Response in Patients with Advanced Melanoma. Clin Cancer Res. 2021;27: 131–140.

26. Farahmand S, Fernandez AI, Ahmed FS, Rimm DL, Chuang JH, Reisenbichler E, et al. Deep learning trained on hematoxylin and eosin tumor region of Interest predicts HER2 status and trastuzumab treatment response in HER2+ breast cancer. Mod Pathol. 2021. doi:10.1038/s41379-021-00911-w

27. Bychkov D, Linder N, Tiulpin A, Kücükel H, Lundin M, Nordling S, et al. Deep learning identifies morphological features in breast cancer predictive of cancer ERBB2 status and trastuzumab treatment efficacy. Sci Rep. 2021;11: 4037.

28. Janowczyk A, Zuo R, Gilmore H, Feldman M, Madabhushi A. HistoQC: An Open-Source Quality Control Tool for Digital Pathology Slides. JCO Clin Cancer Inform. 2019;3: 1–7.

29. Hu Y, Su F, Dong K, Wang X, Zhao X, Jiang Y, et al. Deep learning system for lymph node quantification and metastatic cancer identification from whole-slide pathology images. Gastric Cancer. 2021;24: 868–877.

30. Basavanhally AN, Ganesan S, Agner S, Monaco JP, Feldman MD, Tomaszewski JE, et al. Computerized image-based detection and grading of lymphocytic infiltration in HER2+ breast cancer histopathology. IEEE Trans Biomed Eng. 2010;57: 642–653.

31. Sarker MMK, Makhlouf Y, Craig SG, Humphries MP, Loughrey M, James JA, et al. A Means of Assessing Deep Learning-Based Detection of ICOS Protein Expression in Colon Cancer. Cancers. 2021;13. doi:10.3390/cancers13153825

32. Kleppe A, Skrede O-J, De Raedt S, Liestøl K, Kerr DJ, Danielsen HE. Designing deep learning studies in cancer diagnostics. Nat Rev Cancer. 2021. doi:10.1038/s41568-020-00327-9

33. Sounderajah V, Ashrafian H, Aggarwal R, De Fauw J, Denniston AK, Greaves F, et al. Developing specific reporting guidelines for diagnostic accuracy studies assessing AI interventions: The STARD-AI Steering Group. Nat Med. 2020;26: 807–808.

34. Giovagnoli MR, Ciucciarelli S, Castrichella L, Giansanti D. Artificial Intelligence in Digital Pathology: What Is the Future? Part 2: An Investigation on the Insiders. Healthcare (Basel). 2021;9. doi:10.3390/healthcare9101347

35. Sarwar S, Dent A, Faust K, Richer M, Djuric U, Van Ommeren R, et al. Physician perspectives on integration of artificial intelligence into diagnostic pathology. NPJ Digit Med. 2019;2: 28.

36. Williams BJ, Lee J, Oien KA, Treanor D. Digital pathology access and usage in the UK: results from a national survey on behalf of the National Cancer Research Institute’s CM-Path initiative. J Clin Pathol. 2018;71: 463–466.

37. Eaden J, Mayberry MK, Mayberry JF. Questionnaires: the use and abuse of social survey methods in medical research. Postgrad Med J. 1999;75: 397–400.

38. Kelley K, Clark B, Brown V, Sitzia J. Good practice in the conduct and reporting of survey research. Int J Qual Health Care. 2003;15: 261–266.

39. Pantanowitz L. Digital images and the future of digital pathology. J Pathol Inform. 2010;1. doi:10.4103/2153-3539.68332

40. Hartman DJ, Pantanowitz L, McHugh JS, Piccoli AL, OLeary MJ, Lauro GR. Enterprise Implementation of Digital Pathology: Feasibility, Challenges, and Opportunities. J Digit Imaging. 2017;30: 555–560.

41. Fraggetta F, Garozzo S, Zannoni GF, Pantanowitz L, Rossi ED. Routine digital pathology workflow: The Catania experience. J Pathol Inform. 2017;8: 51.

